# Arctos: Community-driven innovations for managing biodiversity and cultural collections

**DOI:** 10.1101/2023.12.15.571899

**Authors:** Carla Cicero, Michelle S. Koo, Emily Braker, John Abbott, David Bloom, Mariel Campbell, Joseph A. Cook, John R. Demboski, Andrew C. Doll, Lindsey M. Frederick, Angela J. Linn, Teresa J. Mayfield-Meyer, Dusty L. McDonald, Michael W. Nachman, Link E. Olson, Dawn Roberts, Derek S. Sikes, Christopher C. Witt, Elizabeth Wommack

**Author notes:** These authors contributed equally to this work (Carla Cicero, Michelle S. Koo, Emily Braker). Other authors are listed in alphabetical order. Authors are current or former Officers or Board of Director members for the Arctos Consortium.

## Abstract

Museum collections house millions of objects and associated data records that document biological and cultural diversity. In recent decades, digitization efforts have greatly increased accessibility to these data, thereby revolutionizing interdisciplinary studies in evolutionary biology, biogeography, epidemiology, cultural change, and human-mediated environmental impacts. Curators and collection managers can make museum data as accessible as possible to scientists and learners by using a collection management system. However, selecting a system can be a challenging task. Here, we describe Arctos, a community solution for managing and accessing collections data for research and education. Specific goals are to: (1) Describe the core elements of Arctos for a broad audience with respect to the biodiversity informatics principles that enable high quality research; (2) Highlight the unique aspects of Arctos; (3) Illustrate Arctos as a model for supporting and enhancing the Digital Extended Specimen; and (4) Emphasize the role of the Arctos community for improving data discovery and enabling cross-disciplinary, integrative studies within a sustainable governance model. In addition to detailing Arctos as both a community of museum professionals and a collection database platform, we discuss how Arctos achieves its richly annotated data by creating a web of knowledge with deep connections between catalog records and derived or associated data. We also highlight the value of Arctos as an educational resource. Finally, we present a financial model of fiscal sponsorship by a non-profit organization, implemented in 2022, to ensure the long-term success and sustainability of Arctos. We attribute Arctos’ longevity of nearly three decades to its core development principles of standardization, flexibility, interdisciplinarity, and connectivity within a nimble development model for addressing novel needs and information types in response to changing technology, workflows, ethical considerations, and regulations.

## Introduction

Museum collections are veritable treasure troves of objects and associated data that document biological and cultural diversity across spatial and temporal scales. In recent decades, national and global digitization efforts that promote free and open access to those records have unleashed exciting initiatives in both research and education [1–6]. Furthermore, community science efforts aimed at digitizing museum data have shown that entire communities can be engaged in enhancing the scientific value of museum collections [7]. The vast increase in data available through different platforms has revolutionized interdisciplinary studies in evolutionary biology, biogeography, epidemiology, cultural change, and human-mediated environmental impacts [2, 8–10]. Although collections data are increasingly accessible, initiatives for research, education, and policy benefit the most from carefully curated, high-quality information that comprehensively assembles and links everything that is known about objects in an extended network [11–13].

Collection information management systems range from simple spreadsheets to sophisticated relational databases. Fortunately, advances in informatics focused on biodiversity and cultural heritage have enabled broad-scale aggregation of museum data [14] from different sources through the development of global metadata standards (e.g., Darwin Core, https://dwc.tdwg.org; Dublin Core, https://dublincore.org; Getty Vocabularies, https://www.getty.edu/research/tools/vocabularies). Although these efforts have massively increased the *quantity* of data that are available, the *quality* of data depends strongly on local controls that standardize and improve the consistency of data *values* [11]. Efforts to standardize data benefit from community input, especially when diverse disciplines with varying perspectives are represented [15]. Likewise, FAIR Data Principles for scientific data management and stewardship (Findability, Accessibility, Interoperability, and Reusability [16]) promote discovery and use of data through transparency, reproducibility, and reusability.

Museum curators and collection managers are faced with a bewildering number of challenges and choices when considering collections digitization, management, and data access. Although collections data are increasingly available online, not all collection management systems have the advanced infrastructure needed to integrate diverse data sets, broaden the scope of accessible data as new technologies become available, and examine complex interactions and processes [17]. Here, we describe Arctos (https://arctosdb.org), a community solution for managing and accessing collections data for research and education. Specific goals to: (1) Describe the core elements of Arctos for a broad audience with respect to the biodiversity informatics principles that enable high quality research; (2) Highlight the unique aspects of Arctos; (3) Illustrate Arctos as a model for supporting and enhancing the Digital Extended Specimen [12, 18]; and (4) Emphasize the role of the Arctos community for improving data discovery and enabling cross-disciplinary, integrative studies within a sustainable governance model.

### A brief history of Arctos

The foundation of Arctos was set in 1996 when the Museum of Vertebrate Zoology (MVZ) at the University of California Berkeley developed an information management model for its collections (“MVZ Database Model”, [19]). This model was unique at the time in its ability to integrate and manage data from diverse collection types in a single environment, to relate cataloged objects across different collections, and to track and promote access to researchers and educators. The model was implemented as a web-based system and renamed Arctos in 1999 at the University of Alaska Museum (UAM) as part of the Arctic Archival Observatory (National Science Foundation grant DEB-9981915). The University of New Mexico Museum of Southwestern Biology (MSB) and the MVZ began using this Arctos platform in 2003 and 2008, respectively. In 2009, a separate installation of Arctos that eventually became MCZBase was established at the Museum of Comparative Zoology, Harvard University, as a centralized repository for its collections data. The Texas Advanced Computing Center (TACC) began working with Arctos in 2008, first to host media and later to host and provide database support and security for the entire shared system.

Arctos has averaged ∼8-9% annual increase in collection records since its inception over 20 years ago (Fig 1A). Records served by Arctos are globally distributed (Fig 1B), and growth has been concomitant with diversification in the types of data served. Initially developed for vertebrate collections, the University of Alaska Museum became the first institution to add object records from cultural collections in 2014. Arctos now serves rich data across a spectrum of collections beyond vertebrates including archaeology, archives, art, botany, entomology, ethnology, geosciences, history, invertebrate zoology, meteoritics, parasitology, paleontology, teaching, and zooarchaeology. This breadth of data types, along with the accompanying expertise of curators who use Arctos for data management, fosters cross-disciplinary discussions and promotes scientific collaboration and integration.

**Fig 1.**
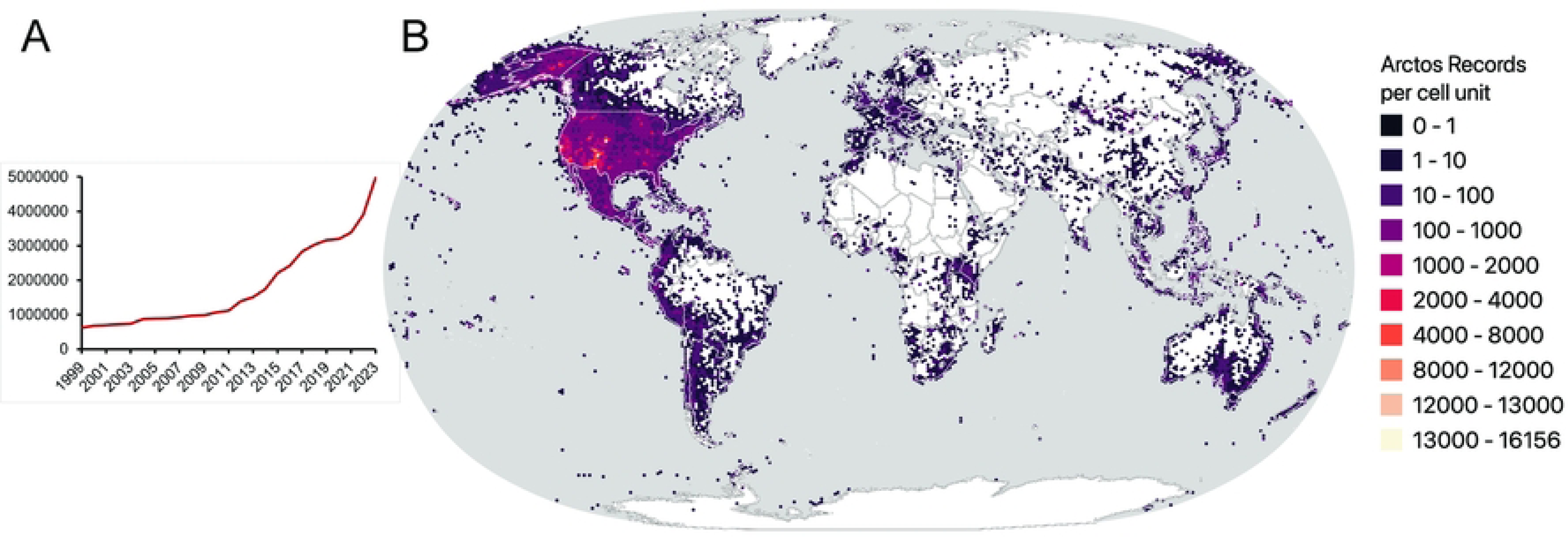
Growth and geographic distribution of Arctos records. (A) Growth of data in the Arctos Consortium showing total number of cataloged records by year (1999 through July 2023). Arctos has grown from ∼614K records in 1999 to ∼5 million records in 2023. (B) Global map of georeferenced localities in Arctos per 100 km2 grid, showing 777,380 spatially distinct localities for over 4 million georeferenced records worldwide with concentrations in North America and Alaska.

### Core features of Arctos

Arctos is a community of museum curators, collection managers, researchers, and informatics professionals as well as a database platform for cutting-edge collection management. As such, it provides a full suite of features for governing, hosting, managing, and connecting collection object data, people, transactions, and other information relevant to collections-based research and education. Arctos is implemented in the relational database PostGRESQL/PostGIS controlled by a Virtual Private Database (VPD, [20]) with a Lucee-based web interface, which allows each collection to manage their data independently. Core versions of its software are released under the open-source license Apache 2.0. (https://github.com/ArctosDB).

As a collection management platform and data portal, Arctos provides a comprehensive solution for managing biological, educational, and cultural collections of all sizes (Fig 2) for museums, universities, state and federal agencies, and field stations. In addition, it functions as its own data aggregator and publisher with dynamic (non-static) object-based data housed in multiple collections and institutions. Because it is a hosted and entirely web-based service, individual collections do not need to spend time or financial resources installing or updating software, maintaining servers, responding to security threats, or coordinating backups. Furthermore, the centralized packaging and publishing of records to external data aggregators (e.g., VertNet, [14]; Global Biodiversity Information Facility [GBIF, 21]) frees collection staff from handling this often-cumbersome process. Arctos is supported through a combination of subscription-based fees, external grants, donations, and in-kind support in the form of personnel subsidized by its members. Subscriptions are on a sliding scale based on collection size and ability to pay, and fee waivers are granted to a small number of collections that lack funding support. The Arctos software development model of “release early, release often” means that it responds quickly as research or collection management needs arise within the consortium. Furthermore, the data model can accommodate a wide variety of data types and values as new collections are added.

**Fig 2.**
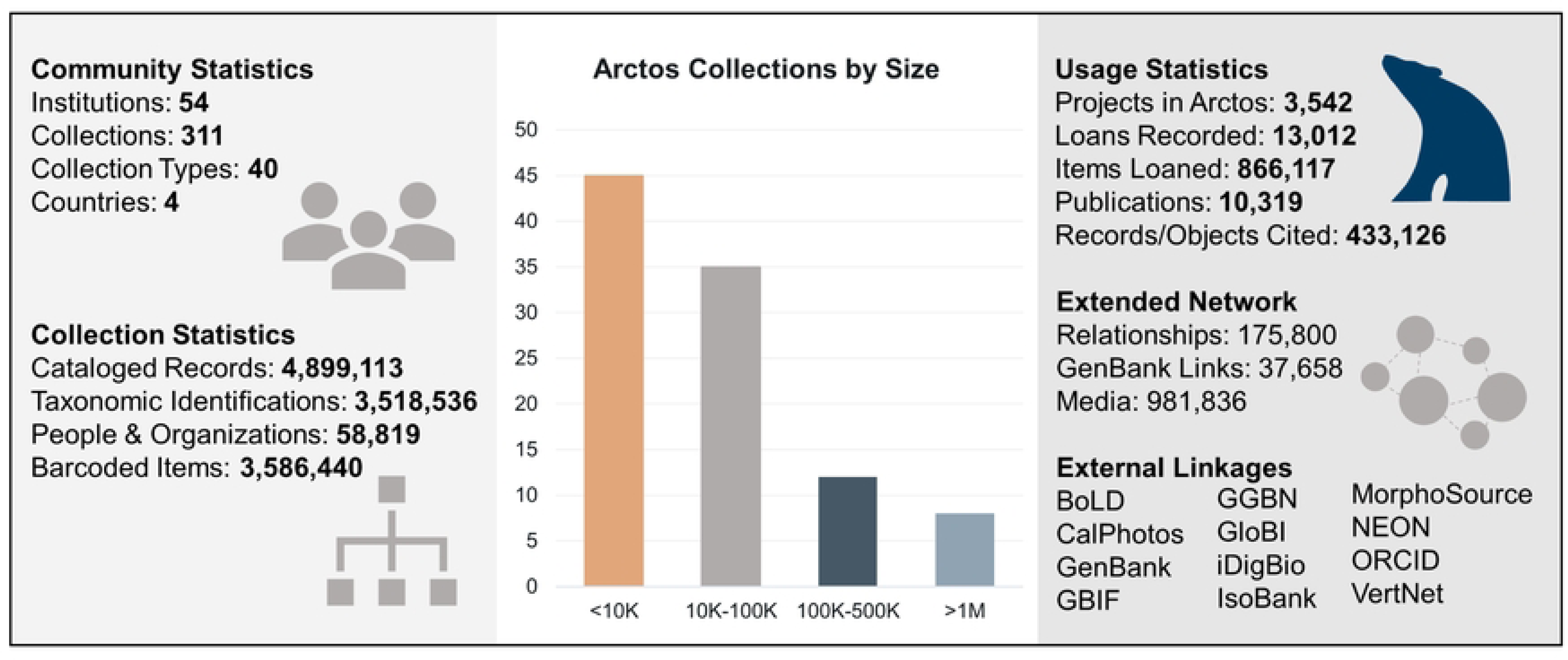
Snapshot of Arctos Collection Management System statistics. Arctos Collection Management System statistics across all collections as of 1 July 2023.

The flexibility of Arctos, combined with its focus on displaying all that is known about a collection object through its integrated data ecosystem, provides a rich platform for scientific and cultural discovery. Its feature-rich components can be categorized into four core areas (Table 1) that are described more fully below: (1) Community; (2) Records; (3) Tools; (4) Connectivity.

**Table 1.**
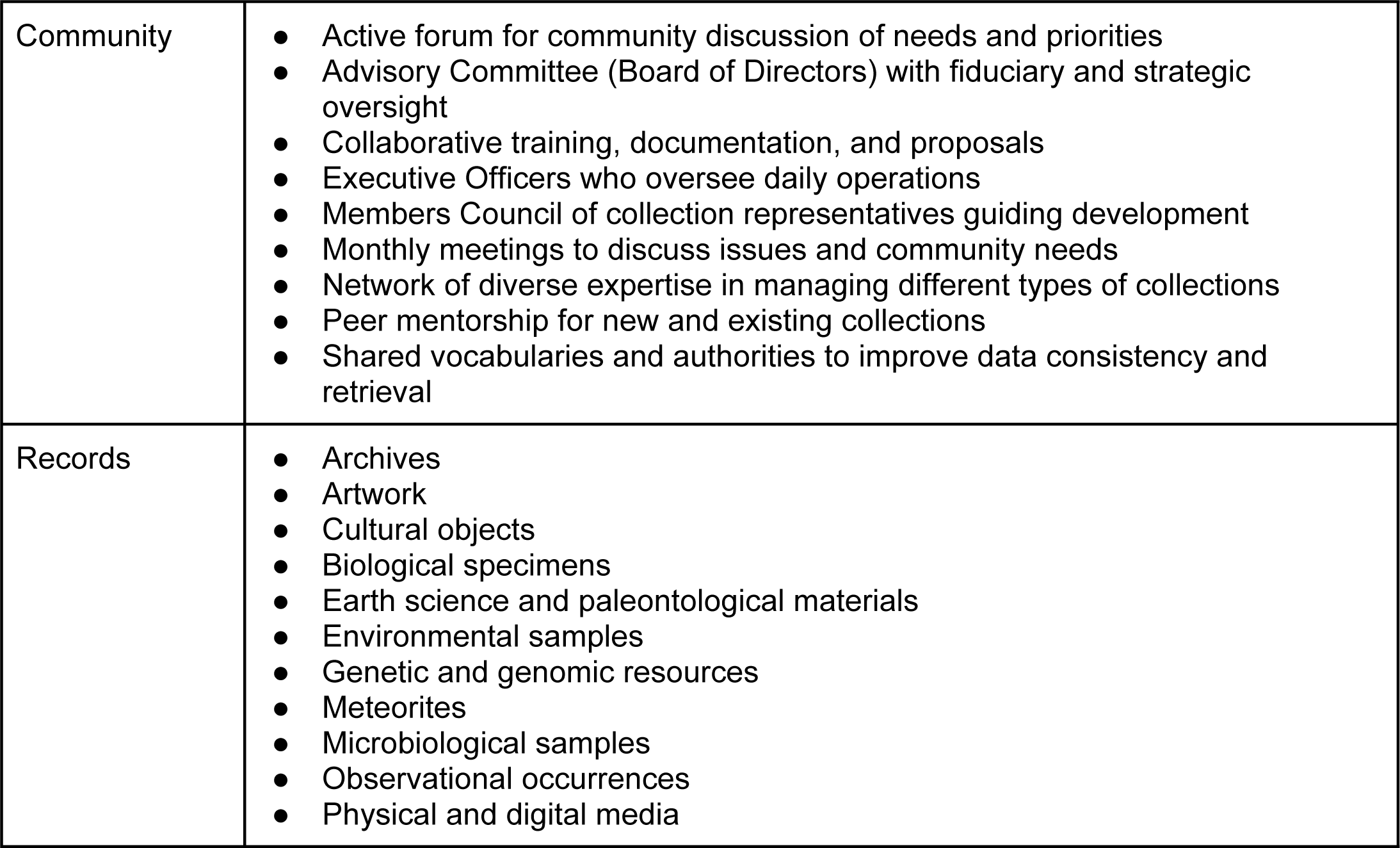
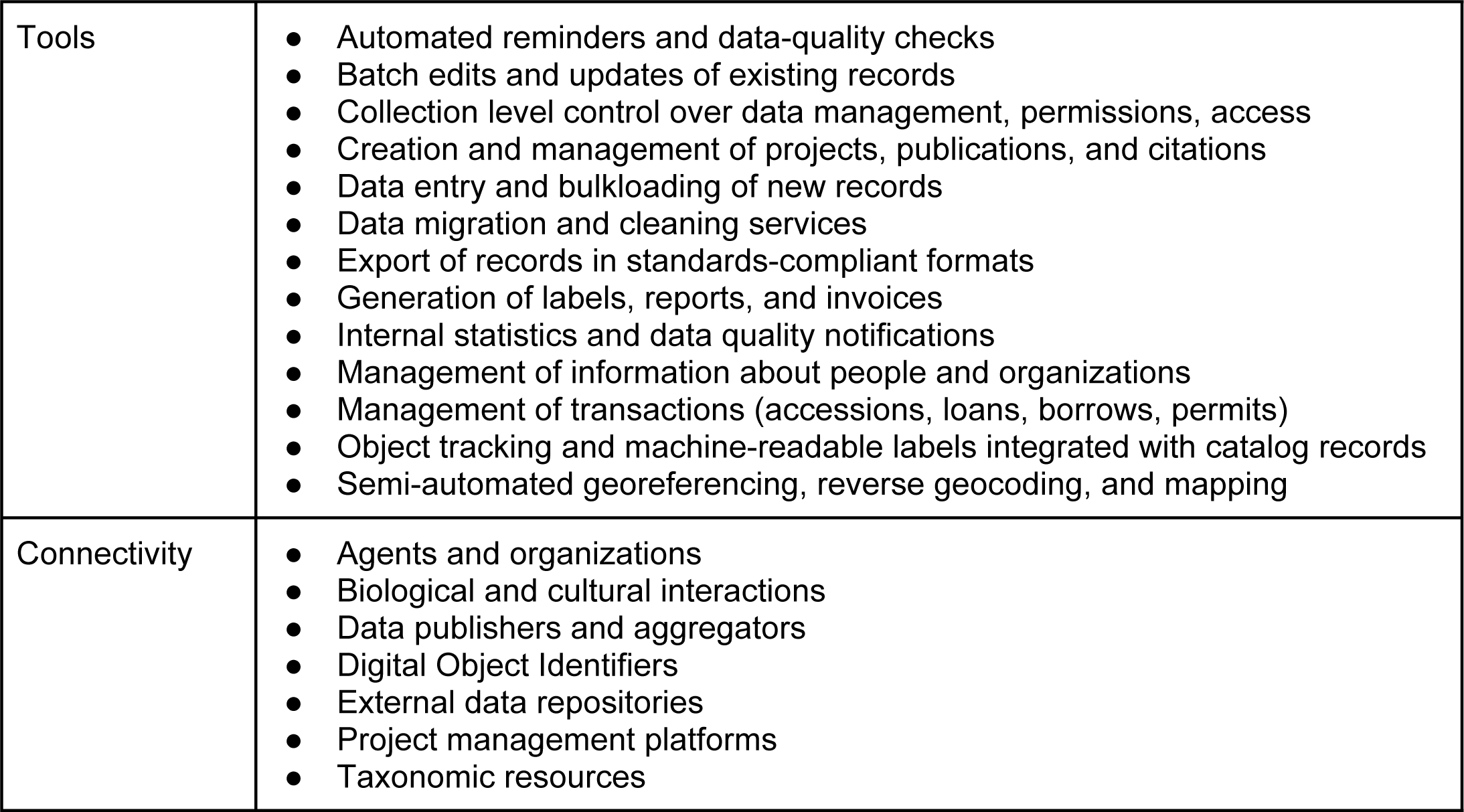
Summary of the core elements of Arctos.

|  |  |
| --- | --- |
| Community | • Active forum for community discussion of needs and priorities<br>• Advisory Committee (Board of Directors) with fiduciary and strategic oversight<br>• Collaborative training, documentation, and proposals<br>• Executive Officers who oversee daily operations<br>• Members Council of collection representatives guiding development<br>• Monthly meetings to discuss issues and community needs<br>• Network of diverse expertise in managing different types of collections<br>• Peer mentorship for new and existing collections<br>• Shared vocabularies and authorities to improve data consistency and retrieval |
| Records | • Archives<br>• Artwork<br>• Cultural objects<br>• Biological specimens<br>• Earth science and paleontological materials<br>• Environmental samples<br>• Genetic and genomic resources<br>• Meteorites<br>• Microbiological samples<br>• Observational occurrences<br>• Physical and digital media |
| Tools | • Automated reminders and data-quality checks<br>• Batch edits and updates of existing records<br>• Collection level control over data management, permissions, access<br>• Creation and management of projects, publications, and citations<br>• Data entry and bulkloading of new records<br>• Data migration and cleaning services<br>• Export of records in standards-compliant formats<br>• Generation of labels, reports, and invoices<br>• Internal statistics and data quality notifications<br>• Management of information about people and organizations<br>• Management of transactions (accessions, loans, borrows, permits)<br>• Object tracking and machine-readable labels integrated with catalog records<br>• Semi-automated georeferencing, reverse geocoding, and mapping |
| Connectivity | • Agents and organizations<br>• Biological and cultural interactions<br>• Data publishers and aggregators<br>• Digital Object Identifiers<br>• External data repositories<br>• Project management platforms<br>• Taxonomic resources |

### Community

The Arctos community is a collaborative, self-governing consortium of collection professionals, information experts, researchers, and educators from diverse disciplines and institutions. As a community, Arctos’ priority is to make research-grade collection data openly accessible and richly networked for multidisciplinary research and public understanding of natural and cultural history. It also serves as a repository and public portal for curated data associated with specimens under federal ownership (e.g., those collected on lands administered by agencies such as the U.S. National Park Service and Bureau of Land Management, among others), and supports or enhances the mission of government agencies at all levels. Community members share in the governance, policy writing, maintenance, and development of Arctos as a collection management platform and data portal (https://arctos.database.museum). In addition to sharing knowledge and expertise, participants form a network of peers that are available to mentor new and existing collection representatives, create training modules (e.g., tutorials, https://arctosdb.org/learn/tutorial-blitz; webinars, https://arctosdb.org/learn/webinars; documentation, https://handbook.arctosdb.org), and discuss development needs and priorities. A shared data environment compels Arctos users to collectively manage controlled vocabularies across different nodes of the database (e.g., geography, taxonomy, agents, preparations, attributes) to promote data standardization and discovery. Consequently, proposed vocabulary or changes to the functionality of Arctos undergo a community decision-making process, ensuring that database developments are guided by Arctos users and reflect community needs. By integrating datasets across biological, geological, and cultural collections, Arctos brings together varied perspectives and data types that lead to innovative, integrative, and broadly beneficial new features and capabilities.

In the collaborative community model, all Arctos collections, regardless of size, are able to actively participate in development priorities and are given equal access to mentors, decision processes, programming aid, community discussion boards, and resources. Learning how to use and contribute to Arctos is a collaborative process which includes regular dialogue among data managers and users. This is especially beneficial to personnel who are less experienced or are interested in growing their knowledge of scientific tools for collection management. Although consensus-building presents its own challenges, the Arctos model encourages community-based solutions, workflow efficiencies, and data quality improvements, thereby advancing best practices in collection data management and data fitness for use [11, 22].

Arctos governance consists of an Advisory Committee and a Working Group composed of volunteer officers, institutional members, technical staff, and subcommittees focused on particular database functions (Fig 3). Regular meetings and online communications through GitHub enable the community to discuss specific issues, address questions or concerns, and resolve problems. Community discussion often focuses on Arctos data that are shared among all collections (e.g., taxonomy, geography, preparations, people, and organizations). Because data standardization is a core tenet of Arctos, database features and enhancements are forged from input and discussion among Arctos users. New features requested by one collection and approved by the community result in a benefit to the community as a whole. Thus, Arctos can be highly responsive to emerging innovations and community needs.

**Fig 3.**
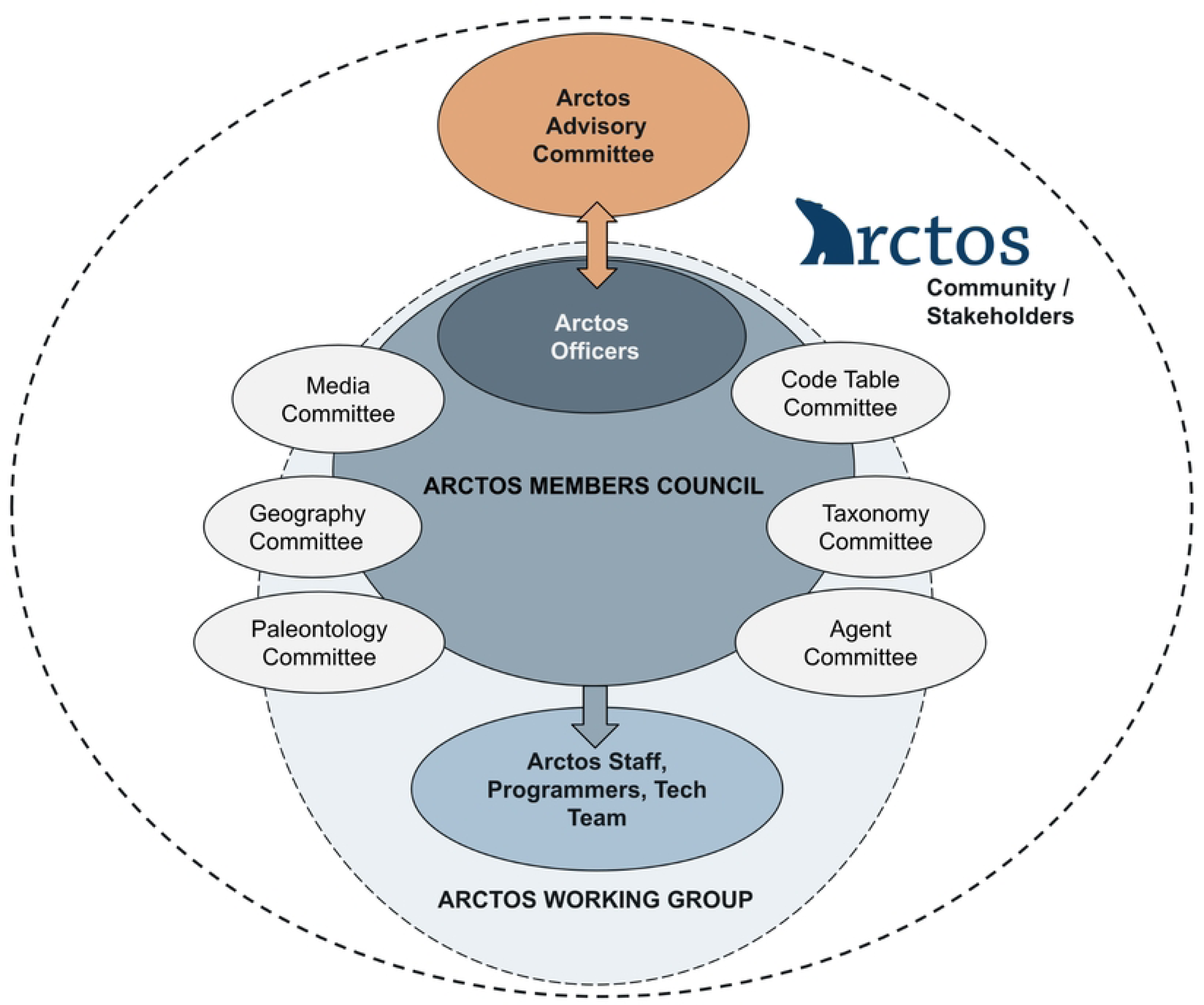
Arctos Consortium organization chart. The Arctos Working Group drives development priorities, responds to issues, engages in outreach, and produces documentation, among other activities. Strategic and financial planning are overseen jointly by the Arctos Advisory Committee and Arctos Officers. Committees within the Working Group focus on specific issues identified by the Arctos community and meet regularly or ad hoc depending on need (see https://arctosdb.org/contacts for details on all Arctos committees).

Shared data lead to positive benefits in efficiencies and data quality improvements. For example, locality georeferences created by one collection are available to other collection records from the same place, forming a gazetteer of vetted data and reducing redundant staff effort (Fig 4). As a case study, the MVZ acquired an orphaned bird collection in 2005 and was able to match over 60% of the records with georeferenced localities already in Arctos. Beyond gained efficiencies, data collated across Arctos institutions enable novel discoveries about people, organizations, and other parties (e.g., businesses, societies, museums, zoos, and government agencies, all of which are included as ‘agents’ in the database) and their contributions within and beyond Arctos (Fig 5). Biographical and statistical information in Arctos produces a holistic view of an individual’s career-long activities across institutions rather than partitioning that information by institution. It also provides opportunities to discover that names associated with different collections are indeed the same person, thus allowing for reconciliation of name variants and corroboration of low-resolution identities while making the data richer and more complete. Finally, the ability to store and share biographical information about agents across collections is an important feature of Arctos, especially for cultural and archival records.

**Fig 4.**
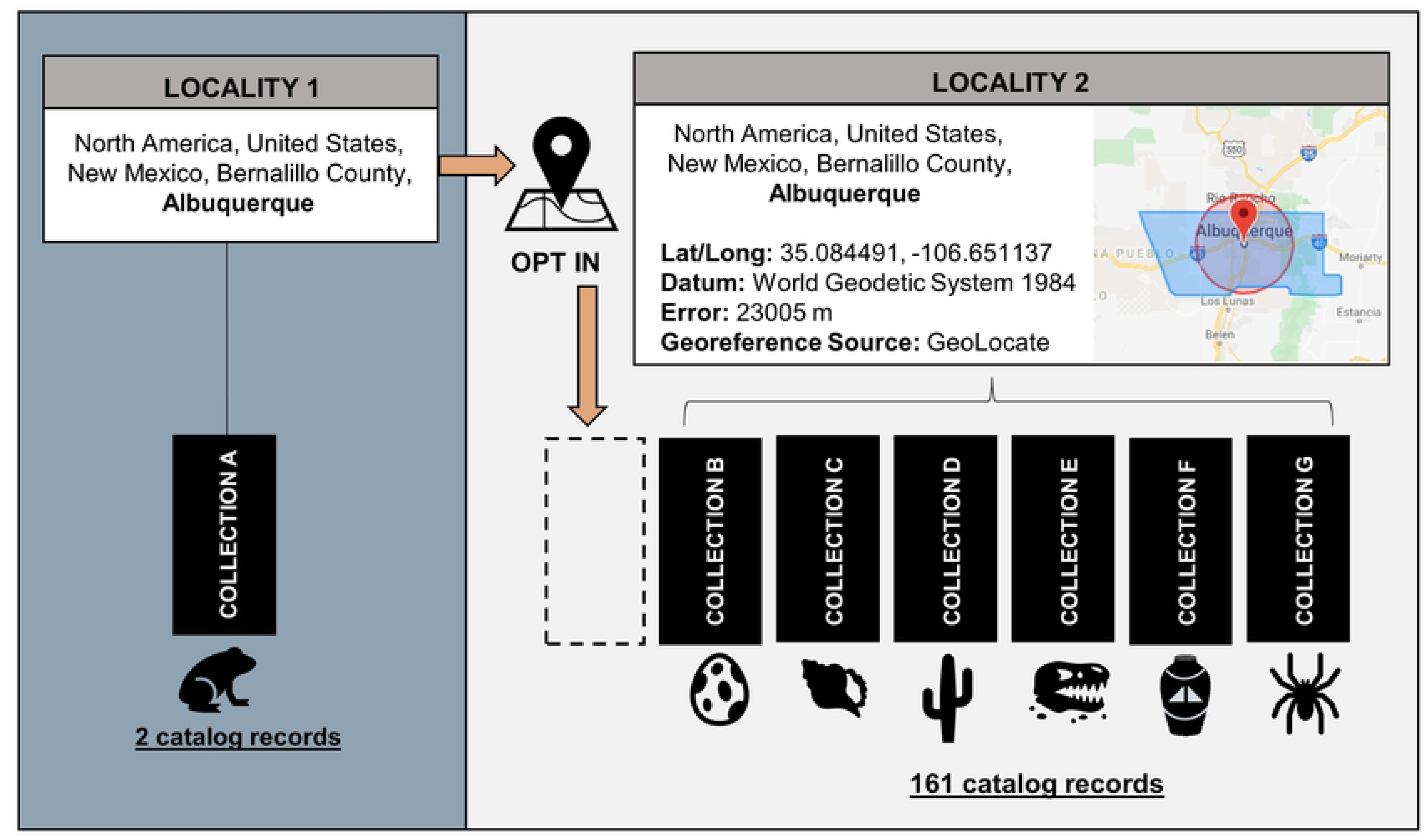
Example of how localities and associated georeferences may be shared in Arctos. Data managers can choose to apply georeferences for a specific locality (e.g., Locality 2) to their own cataloged records that may be from the same descriptive place (e.g., Locality 1) but are lacking coordinates and associated metadata.

**Fig 5.**
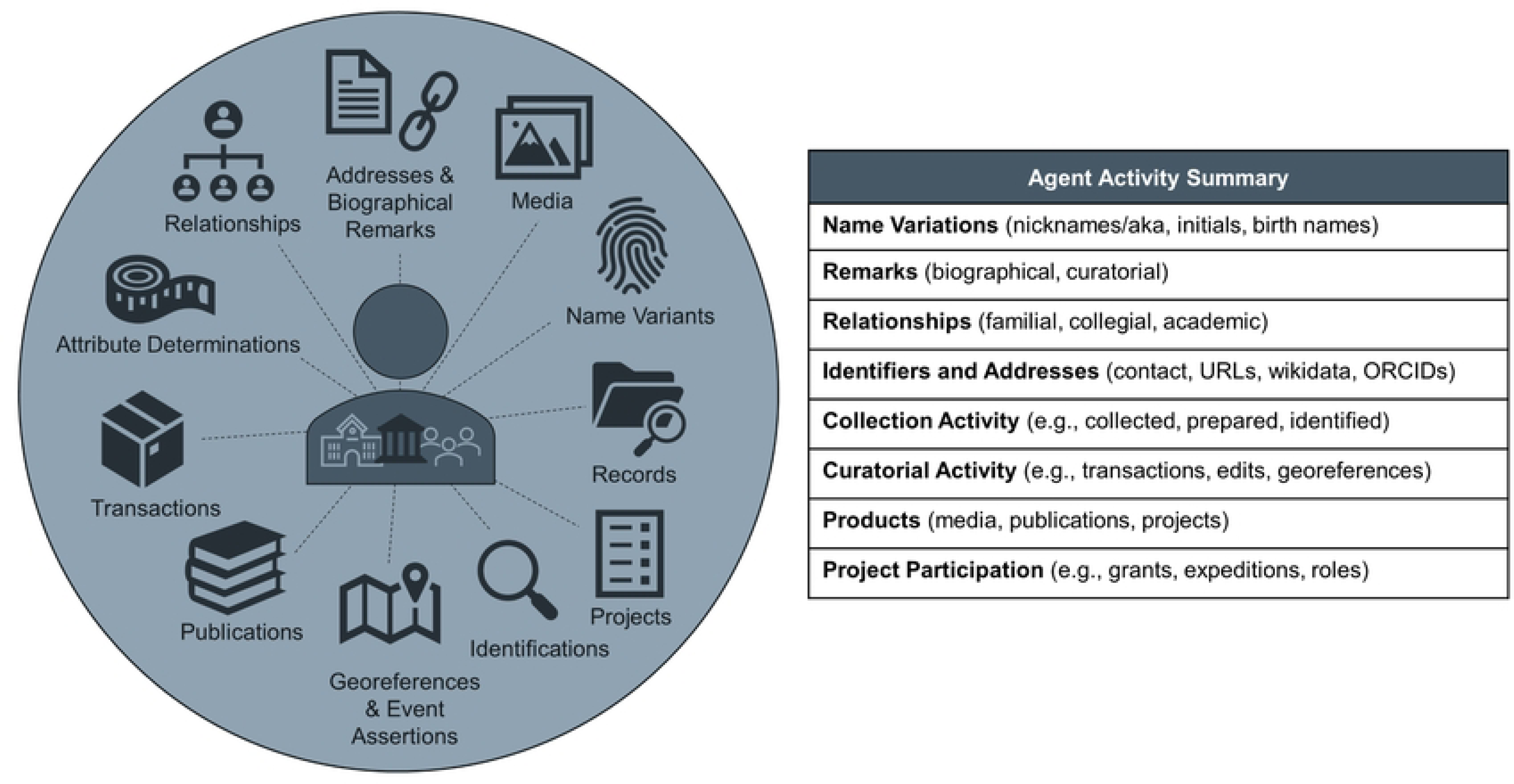
Synopsis of agent activity summary in Arctos. Arctos provides a holistic view of research and curatorial contributions by people, organizations, and institutions (i.e., “agent activity”) to facilitate data relationships, attribution, and assessment metrics. Dynamic links associate agents with related agents (e.g., academic lineages, family members), external resources (wikidata, ORCID), collection activities (objects collected, prepared, or identified), curatorial work (transactions, edits, georeferences), and projects and products (publications, media, grants, expeditions).

### Records

Arctos serves records on over five million specimens, objects, and observations (biological, archival, cultural, geological, and meteorological) curated by participating collections and institutions. The shared database environment allows curators to manage records that cross disciplines (Fig 6), such as objects with both biological and cultural materials or significance, artworks made of iron that intersect with geology, or fossils with mineral taxonomy. Collections can add a mixture of scientific names and/or associated taxa to the identifications of their objects, thus allowing the records to be useful for ethnographic, geologic, and biodiversity research.

**Fig 6.**
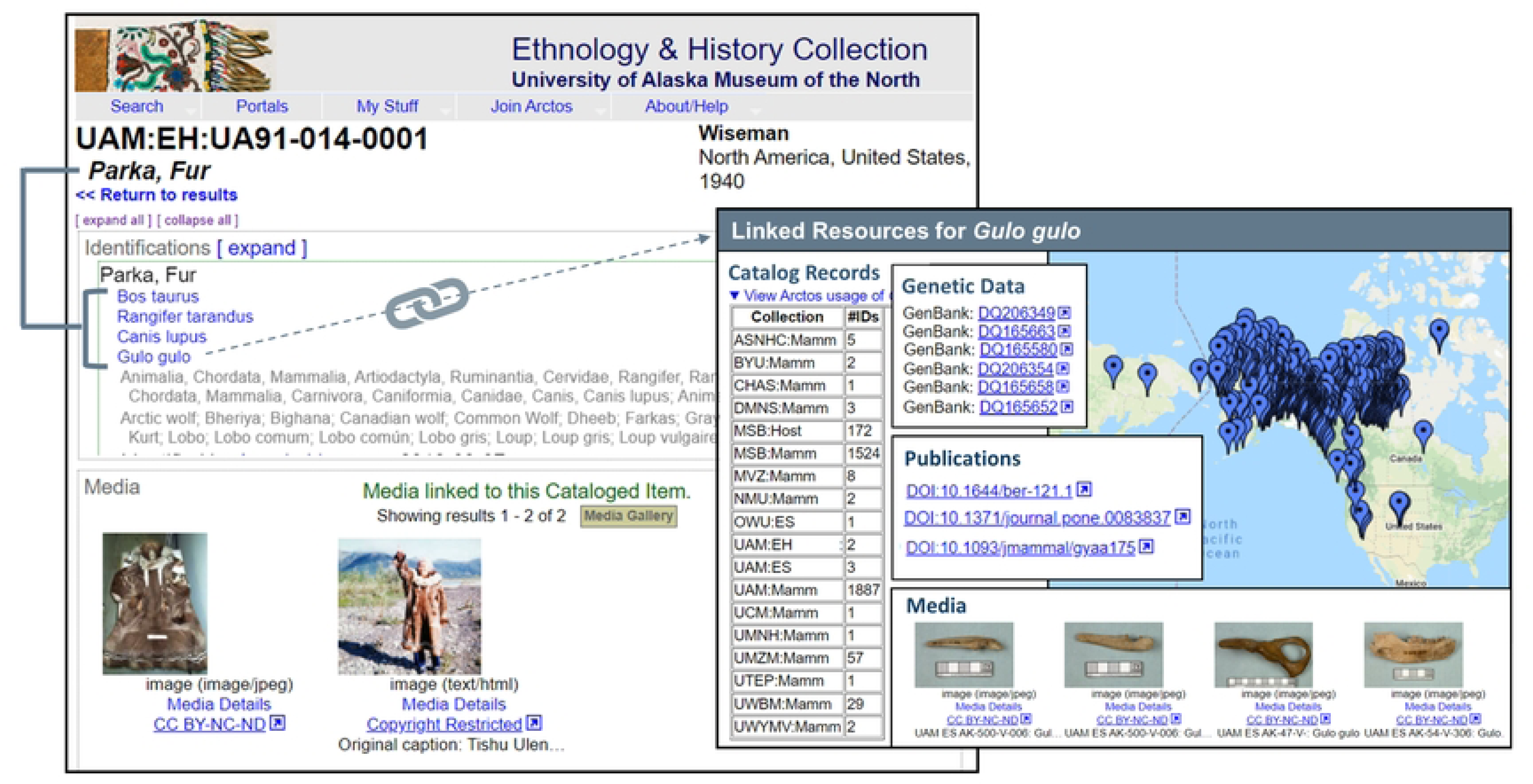
Example of an Arctos record that cross-links cultural and biological records. Arctos record from the University of Alaska Museum Ethnology and History Collection (https://arctos.database.museum/guid/UAM:EH:UA91-014-0001) showcasing a cultural object composed of biological materials that are cross-linked to biological records in Arctos.

Nearly one million media records (e.g., still images, sound recordings, video files) enrich the data in Arctos. Media may showcase objects and specimens before and after preservation [23–24] or function as a voucher for non-specimen observations. In addition, they can be key to increasing access to rare or fragile collection objects that are not typically loaned, such as eggs and nests [25–26], type specimens [27], and objects that are no longer available [28–29]. Increasingly, specimens are being used in high-resolution photogrammetry and three-dimensional scanning projects [30–32] that can be linked directly to the Arctos record. Finally, Arctos media that are linked to collecting events add value by documenting field work, landscapes, habitats, and people.

### Tools

Collection management tools are the nuts and bolts of Arctos functionality, and are used for basic data entry, editing, and searching as well as to improve data quality and increase discoverability. Arctos reports generate transaction invoices, collection ledgers, and dry or wet labels. Different tools find duplicate agents, gaps in catalog numbers, records without parts, unreciprocated relationships between two cataloged items, and records that potentially correspond to GenBank (National Center for Biotechnology, NCBI) accessions but need verification and linking, among other data quality issues. Collection contacts may receive specific reminders with notifications about loans that are due or permits that are expiring. Statistics generated across all of Arctos, or filtered by collection, provide summaries of specific data such as numbers of cataloged items, localities, georeferenced localities, collecting events, agents, media, publications, GenBank links, and specimen relationships.

Access to different tools depends on a user’s training and role. For example, students and volunteers may be given access to enter data for a specific collection by choosing values from existing controlled vocabularies but may not create new taxonomies, higher geography, or names of people and organizations. The addition of new values to controlled vocabularies is limited to collection curators or focused Arctos committees. This hierarchy of permissions limits misspellings and duplication of values (e.g., the same person entered multiple ways), ensures that entries are verified (e.g., a country, state, or county is valid), and compels consistent values for certain fields (e.g., sex, preparations). Ultimately, the focus on standardizing data *values* leads to higher data quality [11] and increases discoverability for researchers and educators using specific criteria.

Below we describe how tools for the following core functions operate in Arctos: data entry and encumbrances; taxonomy and identifications; transactions; object tracking; and spatial data quality.

### Data entry and encumbrances

New records are entered individually or through a variety of batch tools, validated by Arctos through a series of data checks and accepted by curators prior to data loading. Arctos also has implemented workflows to capture data in batches from digitized collection objects such as herbarium sheet labels. Once the data are added, they are immediately accessible online unless a curator chooses to encumber (i.e., restrict access to) those records. Encumbrances may protect sensitive data such as sacred cultural information, collecting locations for fossils or endangered species, and archaeological resources. The Arctos community has developed guidelines for data redaction measures as required by paleontological and cultural collections to meet U.S. federal and private land regulations (e.g., Paleontological Resources Preservation Act of 2009, 16 U.S.C. § 470aaa 1-11; National Historic Preservation Act of 1966, Public Law 89-665; 54 U.S.C. 300101 *et seq;* and Archaeological Resources Protection Act of 1979, 16 U.S.C. 470aa-470mm; Public Law 96-95 and amendments). Likewise, encumbrances restrict usage of collection objects and data according to permit conditions and material transfer agreements (e.g. Nagoya Protocol [33]).

### Taxonomy and identifications

Identifications within Arctos are treated separately from taxonomy, both during data entry and editing. Taxonomy refers to formal classification systems including cultural lexicons, and Arctos harvests data from external web services (e.g., Global Names Architecture, https://globalnames.org) while allowing for customized taxonomies. This provides both data quality control and the flexibility of collections to choose and modify their own taxonomy. Collection staff then apply those names to collection objects through a flexible identification module that allows for vernacular, regional, and Indigenous names, taxonomic uncertainties, biological realities (e.g., hybrids, intergrades, new species with temporary designations), multiple identifications, non-hierarchical identifications, and nomenclature from geological, archival, art, and cultural collections. For example, a hybrid specimen is identified by selecting two parental names from the Arctos taxonomy table, and a parka made of furs from different species is identified by multiple taxonomic names that comprise the components (e.g., parka, *Bos taurus, Rangifer tarandus*, *Canis lupus*, *Gulo gulo*; Fig 6). This can also be applied to egg sets with nest parasites, which are identified by the taxonomic name of both the host and the parasite (e.g., *Melospiza melodia* and *Molothrus ater*, https://arctos.database.museum/guid/MVZ:Egg:609). Importantly, Arctos records the history of all changes to identifications; when new identifications and associated metadata (e.g., determiner, date, basis of determination) are added, old but invalid (i.e., erroneous or synonymous) identifications are retained and remain searchable. Arctos also allows batch identification updates, critical for management of entomology as well as other collections.

### Transactions

Transactions include accessions, loans (outgoing collection material), borrows (incoming material from another collection), and permits, all of which are managed in Arctos. Accessions, loans, and borrows are collection-specific, with options for formatting transaction numbers depending on in-house curatorial practices. Permit metadata, on the other hand, are shared among collections and can be linked to accessions, loans, and borrows (the permits themselves are private unless a collection chooses otherwise). This facilitates the management of permits, material transfer agreements, memoranda of understanding, and other documentation for compliance with state, federal, and international (e.g., CITES, Nagoya) regulations [33]. It also allows specimens or objects acquired by one collection under its permit(s) to be accessioned into a different collection using the same permit(s). Likewise, loans sent by different collections can use the same institutional permit. In addition to capturing basic information about the transaction (i.e., persons and/or agencies involved, date, transaction number, nature and amount of material, remarks), transactions can be linked to media such as images or documents (including spreadsheets) that provide supporting information. Arctos also tracks the use of data *about* objects (e.g., metadata or media) through transactions such as data loans and media loans; users may request data records or object photographs rather than objects per se, allowing more comprehensive documentation about how records, information, and associated elements are being used.

### Projects and publications

A special feature of Arctos is the ability to link transactions to thematic web pages called “Projects” that summarize the contributions and use of cataloged records for activities, studies, or uses (e.g., field trips, digitization initiatives, traveling exhibitions, collections on state or federal lands). Projects also are used to document biosampling contributions by Indigenous communities for resource policy and decision-making as well as to highlight cultural object acquisitions, displays, and practices. Publications and media resulting from these activities are easily added to the project page. Further, publications may be cross-referenced to digital object identifiers (DOIs) and cataloged records may be added as citations, directly showing usage of individual records within or across collections. Thus, projects serve as the central hub that link all related data in Arctos to showcase the impact of collections by highlighting how researchers, educators, organizations, and others are using the platform for discipline-specific or interdisciplinary goals (Fig 7; examples in Table 2). Projects are automatically related to other projects based on shared objects, which enables deep-tracking of a collections’ utility and products through time. They serve as a convenient hub for agencies, funders, and users to access up-to-date records associated with specific activities.

**Fig 7.**
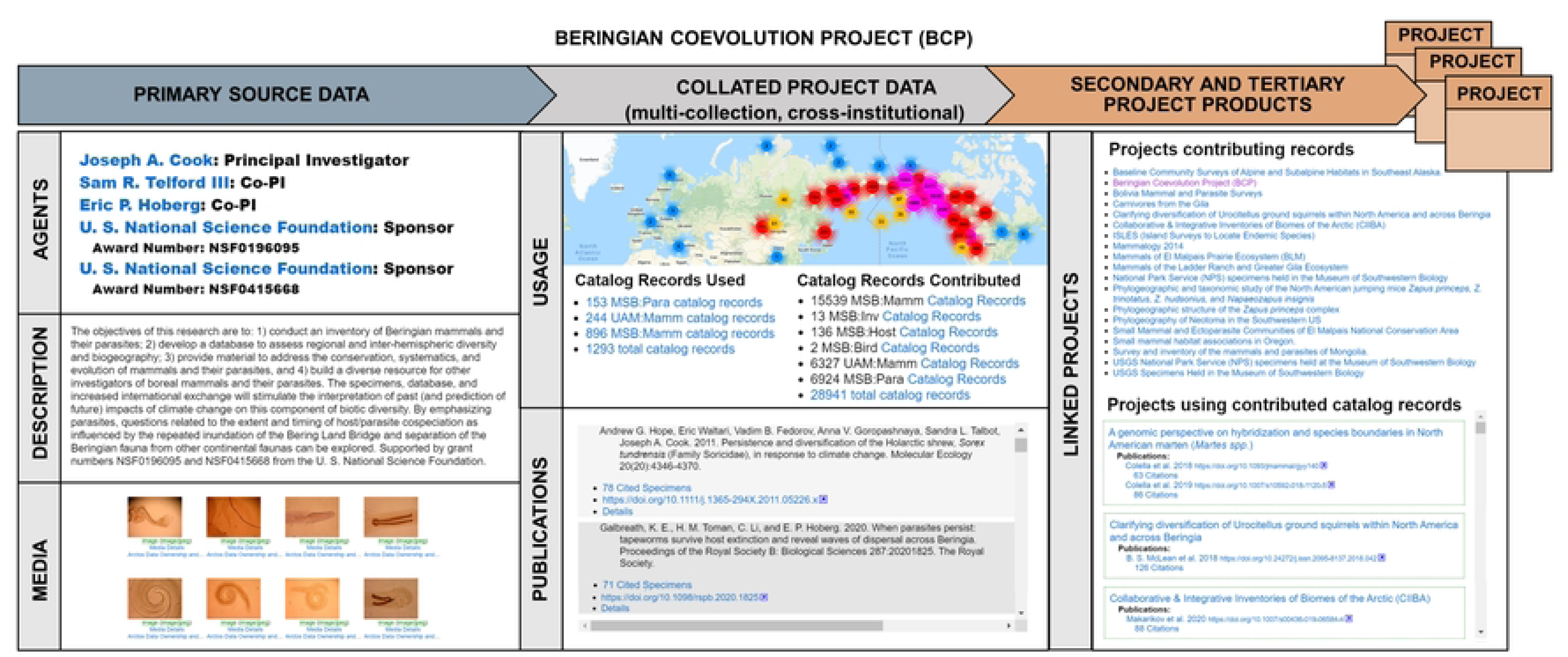
Synopsis of the Beringian Coevolution Project in Arctos. The Beringian Coevolution Project (https://arctos.database.museum/project/51) showcases primary source data and collated products. The project included 28,941 cataloged records representing co-examined boreal mammals and associated parasites from two museums and resulted in 8 media objects, 216 publications citing 8,561 specimens, and a dynamic web of 153 related projects that either contributed records used by this project or that used specimens from this project to generate additional research outputs (293 subsequent publications with 69,087 citations).

**Table 2.** Examples of projects in Arctos that target different users. Arctos project IDs have the base URL https://arctos.database.museum/project/ (e.g., https://arctos.database.museum/project/10000850).

| Topic | Arctos Project ID | Target User(s) |
| --- | --- | --- |
| Alaskan Insect Pollinators | [10000850](https://arctos.database.museum/project/10000850) | Alaska Department of Fish and Game, Researchers, National Science Foundation |
| Alaskan Plant Survey | [1000054](https://arctos.database.museum/project/1000054) | National Park Service |
| Albuquerque BioPark & Museum of Southwestern Biology Specimen Repository Agreement | [10002948](https://arctos.database.museum/project/10002948) | Researchers, Zoos |
| Art and Ethnology Collections purchased with funds from Rasmuson Foundation | [10003033](https://arctos.database.museum/project/10003033) | Artists, Funder |
| Art Work Inspired by Life | [10002855](https://arctos.database.museum/project/10002855) | Artists |
| Beringian Mammal and Parasite Coevolution | 51 (also see Fig. 9) | Researchers, National Science Foundation |
| Center for Disease Control<br>Hantavirus Survey in National Parks | 10002373 | U.S. Center for Disease Control,<br>Epidemiologists, Researchers,<br>National Park Service |
| Educational Collaboration between<br>Art and Biology | 10003671 | Students |
| Support for Ornithology and<br>Herpetology Collections | 10003135 | National Science Foundation |
| Mexican Wolf Recovery Program | 1000071 | U. S. Fish and Wildlife Service,<br>Conservation Groups |
| Resurvey of Vertebrate Communities<br>in California | 10000047 | Researchers, Agencies, National<br>Science Foundation |
| Seal Specimens Hunted by Native<br>Americans in Alaska | 15 | Alaska Native Harbor Seal<br>Commission |
| Greenwood Wildlife Rehabilitation<br>Center Salvaged Vertebrates | 10004199 | Agencies/governments (city,<br>county, state, etc.), Researchers |

### Object tracking

Object tracking in Arctos acts as an independent module linked via barcodes to the catalog record and provides the capacity to track materials from collection and accession through cataloging, storage, and loans. For example, genetic resource collections can be organized in a hierarchy that associates barcoded cryovials with specific positions in barcoded boxes, racks, freezers, and rooms within buildings (e.g., nested “containers”; Fig 8). Similarly, barcodes are used to track the locations of dry specimens in cabinets or fluid-preserved specimens in jars on shelves. Arctos can accommodate different types of scannable codes (e.g., a true barcode) or non-scannable codes (e.g., cabinet numbers printed on labels) that are captured through the container module, and these can be applied to both cataloged and non-cataloged (in process) items. In addition to tracking the location of collection objects, container environments (e.g., relative humidity, temperature, ethanol concentration, Integrated Pest Management check) and their history can be tracked to better monitor collections and document changes over time. Containers in Arctos are broadly applicable to a variety of curatorial functions, including management of accessions and loans, collection inventories and moves, conservation and preservation of collection objects, and Integrated Pest Management practices.

**Fig 8.**
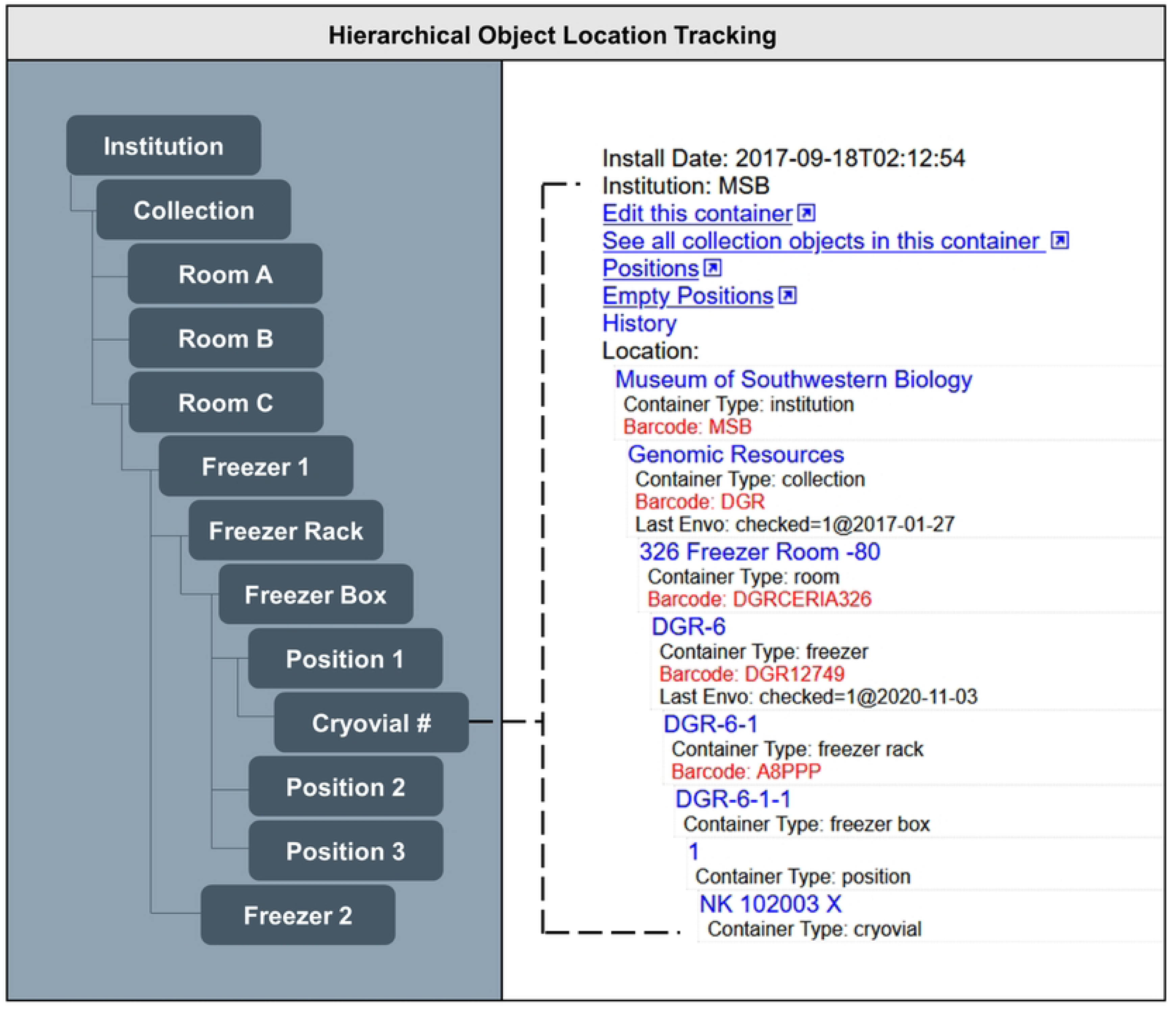
Schematic showing hierarchical object tracking of tissues in Arctos. In this example, the hierarchy shows where an institutional collection freezer is located and the nested position of a cryogenic vial within a box in that freezer.

### Spatial data quality

Geography in Arctos adds critical value to records as fundamental metadata and as a measure of data quality. Adding spatial data allows records to be correlated with environmental and geographic data, thus ensuring their usefulness beyond Arctos [34–36]. Over 75% of the more than 850,000 localities in Arctos have associated georeferences that are available to Arctos users and global data aggregators. Localities are shared among all collections in Arctos, which brings advantages (Fig 4). For example, diverse taxa from the same expedition (e.g., snails, fish, and salamanders collected in the same pond) or collected at different times from the same location can share the same locality in Arctos. This saves georeferencing effort, ensures data consistency, improves discoverability, and stimulates cross-disciplinary integration.

With an emphasis on spatial accessibility and quality, Arctos has a suite of tools for mapping and describing spatial locations of collection objects. A plug-in for GeoLocate (https://geo-locate.org) facilitates semi-automated georeferencing while BerkeleyMapper (https://berkeleymapper.berkeley.edu) provides data visualization and spatial exploration tools. Arctos also uses Google Maps web services for automated data-quality checking, whereby reverse geocoding verifies if coordinates are in the correct higher geography (i.e., continent, country, state/province, county). Higher geographies are defined with polygons, and countries follow a spatially explicit authority (Database of Global Administrative Areas, https://gadm.org) which uses ISO standards (International Organization for Standards, https://iso.org). Polygons are not limited to higher geography but can be used to describe an object’s locality instead of point coordinates. Finally, locality attributes add descriptive terms to a place (see https://handbook.arctosdb.org/documentation/geology.html), and localities can be verified and locked once checked by collectors or curators to preserve data integrity.

### Connectivity

Arctos prizes connectivity, in which everything that is known about an object and its relationships, interactions, or derivatives can be displayed or made accessible. For this reason, Arctos integrates with a growing list of external data repositories and services (Table 3) that add value to its data records. This core feature makes Arctos a uniquely rich center of collection-related data and tools for the exploration and visualization of biological, geological, and cultural diversity in novel ways. For example, Arctos can integrate with any resolvable identifier, and was the first collection management system to develop reciprocal, dynamic connections between specimen records and genetic data in GenBank. Dynamic linkages from GenBank back to the Arctos record are created when submissions to NCBI involve referencing the specimen voucher in the NCBI “specimen_voucher” field using three-part “Darwin Core Triplets” (institution:collection:catalog number). As of July 2023, over 37,600 Arctos records are linked to associated data in GenBank.

**Table 3.**
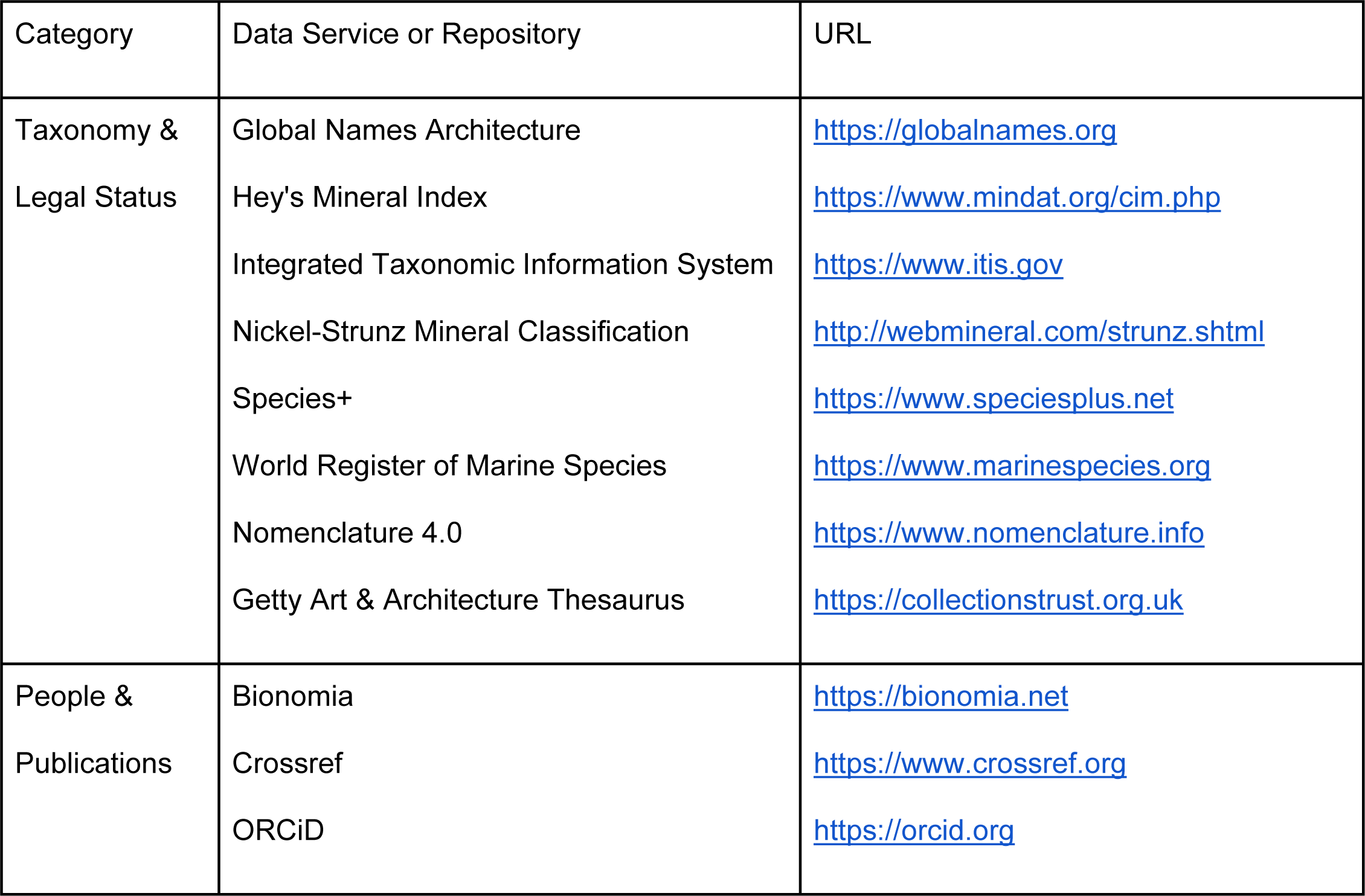
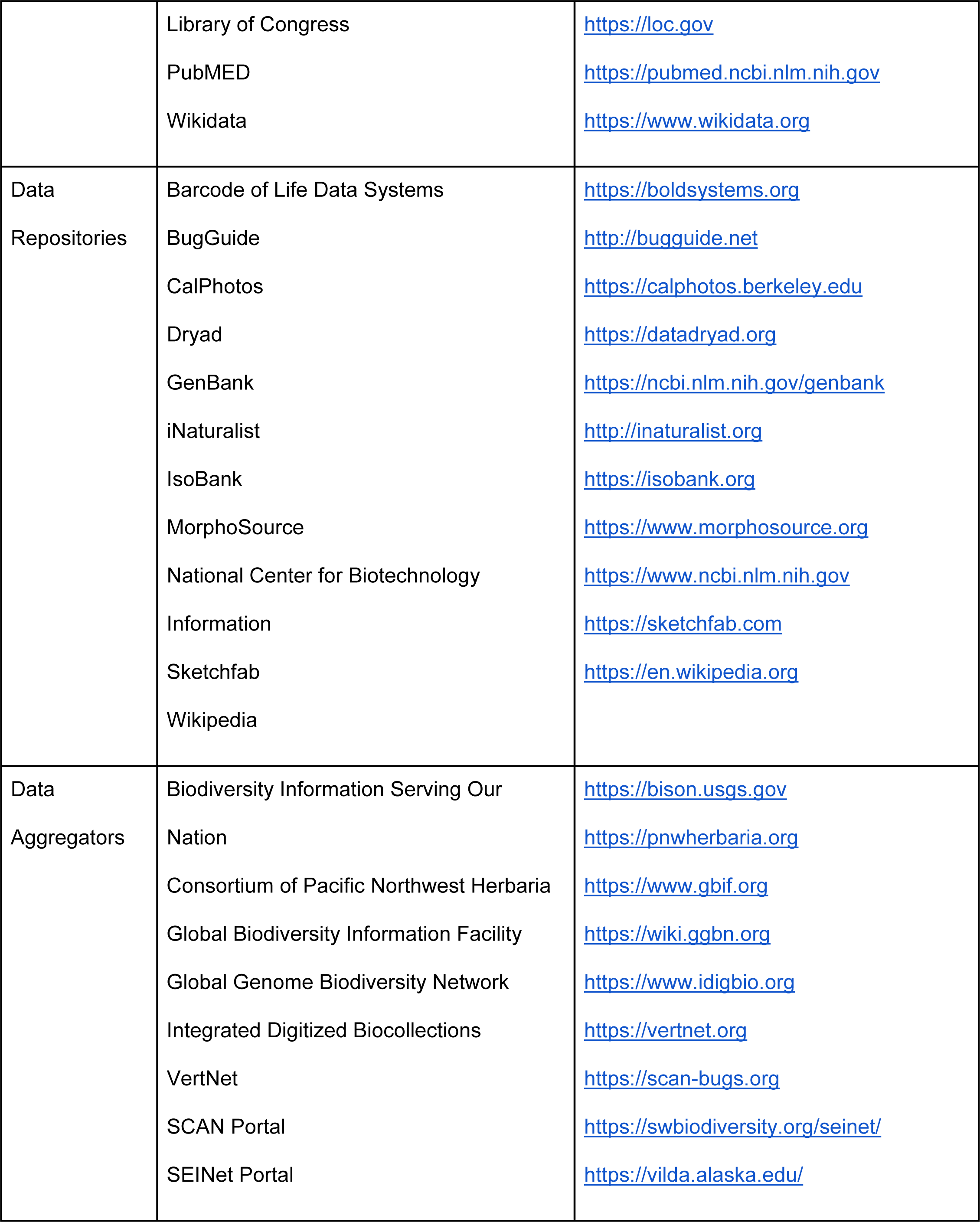
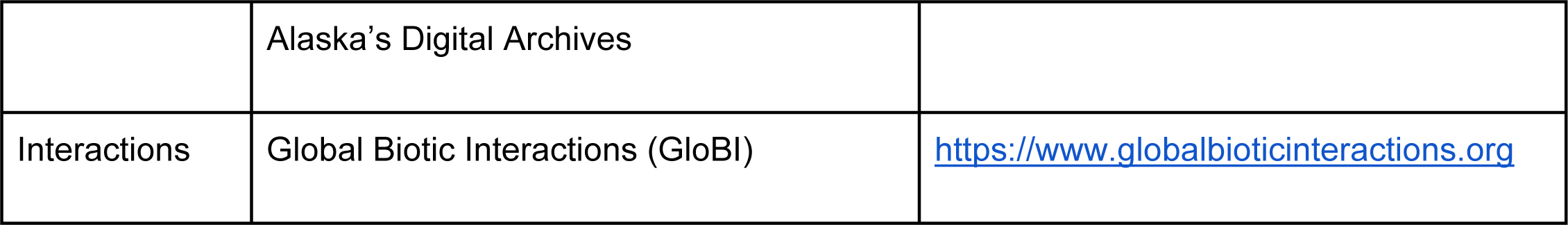
Arctos connections with external repositories, authorities, and databases. Links to these sites add value and relevance for collection transactions, curation, and record management for both natural and cultural history collections. Some connections are reciprocal while others integrate values from an authority. Connections may be made through web services automatically or updated manually.

| Category | Data Service or Repository | URL |
| --- | --- | --- |
| Taxonomy &<br>Legal Status | Global Names Architecture<br>Hey's Mineral Index<br>Integrated Taxonomic Information System<br>Nickel-Strunz Mineral Classification<br>Species+<br>World Register of Marine Species<br>Nomenclature 4.0<br>Getty Art & Architecture Thesaurus | https://globalnames.org<br>https://www.mindat.org/cim.php<br>https://www.itis.gov<br>http://webmineral.com/strunz.shtml<br>https://www.speciesplus.net<br>https://www.marinespecies.org<br>https://www.nomenclature.info<br>https://collectionstrust.org.uk |
| People &<br>Publications | Bionomia<br>Crossref<br>ORCID | https://bionomia.net<br>https://www.crossref.org<br>https://orcid.org |
|  | Library of Congress<br>PubMed<br>Wikidata | https://loc.gov<br>https://pubmed.ncbi.nlm.nih.gov<br>https://www.wikidata.org |
| Data<br>Repositories | Barcode of Life Data Systems<br>BugGuide<br>CalPhotos<br>Dryad<br>GenBank<br>iNaturalist<br>IsoBank<br>MorphoSource<br>National Center for Biotechnology<br>Information<br>Sketchfab<br>Wikipedia | https://boldsystems.org<br>http://bugguide.net<br>https://calphotos.berkeley.edu<br>https://datadryad.org<br>https://ncbi.nlm.nih.gov/genbank<br>http://inaturalist.org<br>https://isobank.org<br>https://www.morphosource.org<br>https://www.ncbi.nlm.nih.gov<br>https://sketchfab.com<br>https://en.wikipedia.org |
| Data<br>Aggregators | Biodiversity Information Serving Our<br>Nation<br>Consortium of Pacific Northwest Herbaria<br>Global Biodiversity Information Facility<br>Global Genome Biodiversity Network<br>Integrated Digitized Biocollections<br>VertNet<br>SCAN Portal<br>SEINet Portal | https://bison.usgs.gov<br>https://pnwherbaria.org<br>https://www.gbif.org<br>https://wiki.ggbn.org<br>https://www.idigbio.org<br>https://vertnet.org<br>https://scan-bugs.org<br>https://swbiodiversity.org/seinet/<br>https://vilda.alaska.edu/ |
|  | Alaska's Digital Archives |  |
| Interactions | Global Biotic Interactions (GloBI) | https://www.globalbioticinteractions.org |

### A model for the multidimensional Digital Extended Specimen

Connectivity is a core principle of the Digital Extended Specimen network, which promotes interdisciplinary research into functional traits [37] and biological interactions [38], provides a critical foundation for global conservation efforts [38], and reflects the role that “next-generation collections” [17] play in advancing science and society. From its inception, Arctos as a data platform has been built on the “extended specimen network” concept - that is, linking physical objects to all of their derived data (especially web-accessible digital assets) and to third-party repositories for increased accessibility and discoverability [13, 18, 39].

Arctos achieves its richly annotated data by creating a web of knowledge with deep connections between catalog records and derived or associated data, and by using reliable published resources for globally shared information. Here we illustrate how the extended specimen in Arctos can become multidimensional (Fig 9). An Alpine Chipmunk (*Tamias alpinus*) was collected with a pinworm parasite (*Rauschtineria eutamii*) in Inyo County, California, in 2010, and the two specimens are accessioned in the Museum of Vertebrate Zoology (https://arctos.database.museum/guid/MVZ:Mamm:225308) and Museum of Southwestern Biology (http://arctos.database.museum/guid/MSB:Para:27057), respectively. Each specimen has its own extended data network with URL-based links to GenBank sequences, media, and a shared georeferenced collecting event. Within Arctos, the extended specimen networks are multiplied by several inter-collection connections: (1) the two specimens are explicitly related to each other with a biotic interaction of host and parasite, and these relationships are harvested by the Global Biotic Interactions platform [40]; (2) they were cited in publications [41, 42] shared across collections; and (3) they share a collecting event with other specimens that may be important to ecological studies of parasites. Lastly, the chipmunk was collected as part of a state-wide effort to resurvey California biodiversity (Arctos project https://arctos.database.museum/project/10000244), placing occurrences for both the mammal and its associated parasite in a research context.

**Fig 9.**
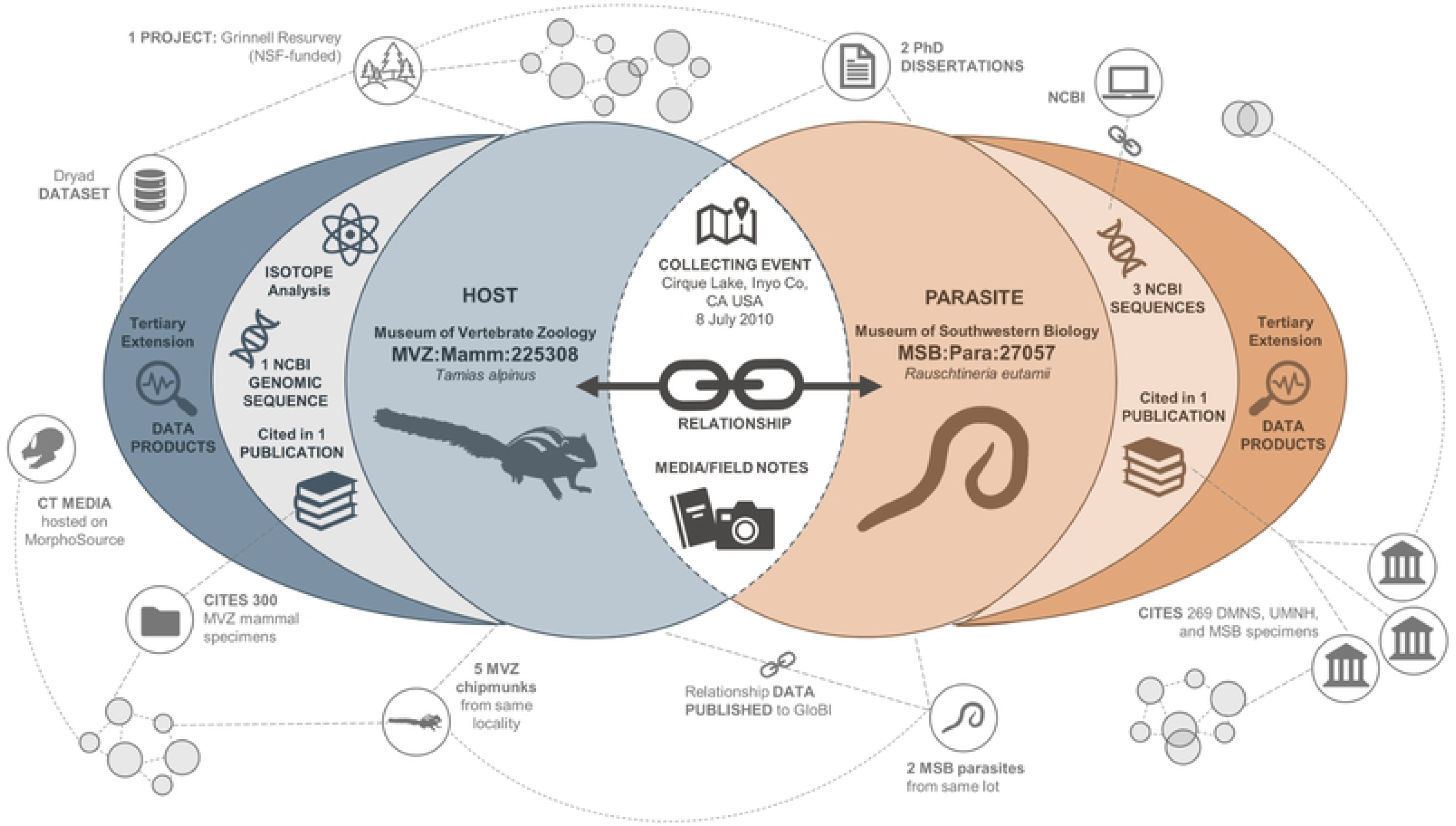
Multidimensional extended specimens in Arctos. Records in Arctos include multidimensional extended specimens that share primary, secondary, and tertiary components and are directly related to each other. The example shown here includes a chipmunk (MVZ:Mamm:225308) and its parasite (MSB:Para:27057) that were collected and accessioned at separate Arctos institutions. They share host-parasite biotic interactions, the same collecting event, and primary source material such as field notes. Any updates to host or parasite metadata (e.g., identification, locality, date) are reflected and searchable in both records across institutions. Together, the chipmunk and parasite were used in graduate student research producing at least two publications and two dissertations that cited 569 specimens from four Arctos collections. The host is linked to one genomic sequence deposited at the National Center for Biotechnology Information (NCBI) Sequence Read Archive (SRA), and the parasite is linked dynamically to three GenBank sequences at NCBI. In addition, the chipmunk was one of 10,987 vertebrate specimens and observations collected in the southern Sierra Nevada as part of the Grinnell Resurvey Project [41], which resulted in 19 more publications and 2,215 citations in Arctos.

### The Arctos Entity

The premise of the Digital Extended Specimen revolves around an individual specimen or object with a single collecting event and links to its derivatives such as gene sequences, CT scans, isotope data, and media (Fig 9). However, the reality of many collections may be more complicated. An individual specimen may be split or composed of separate components, each of which may have different collecting events, preparators, preservation types, and other metadata. Arctos community discussion on how to address this challenge led to the creation of a network-wide “Entity” collection that acts to combine multiple component records sharing the same organism, object, or event ID into a single dashboard with a unique, shareable URL (Fig 10). These components are linked from the Arctos entity record to their own respective record via individual URLs.

**Fig 10.**
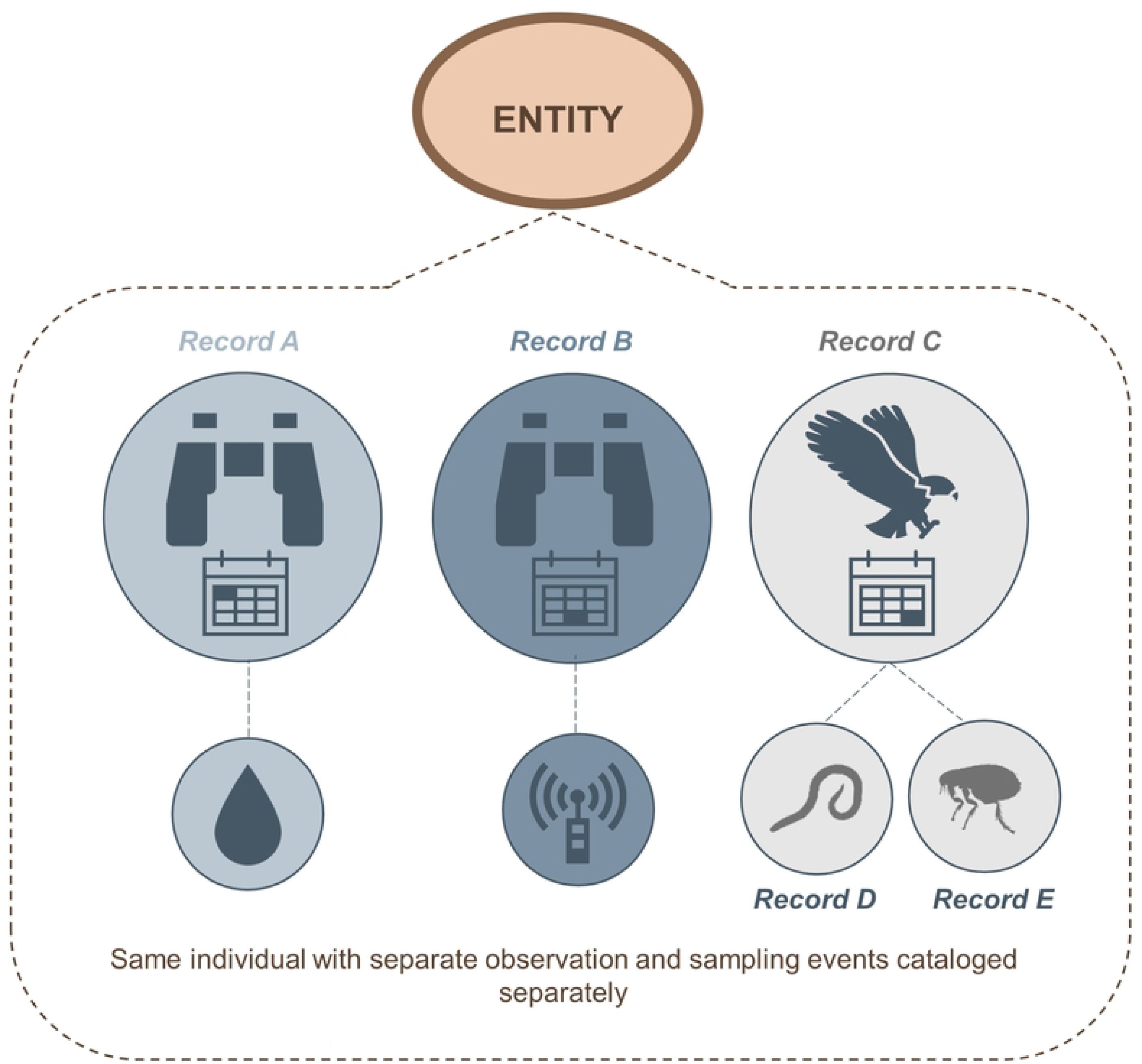
Example of the Arctos Entity model. The Arctos Entity model links diverse records to a single unifying record for increased discoverability. Here, Record A is a bird observation with an associated blood sample, Record B is a second observation of the same individual with associated radio telemetry data, Record C is the vouchered specimen, and Records D and E are endoparasites and ectoparasites, respectively, taken from the specimen.

For example, one entity record (https://arctos.database.museum/guid/Arctos:Entity:16) links the cataloged blood sample of a Golden Eagle (*Aquila chrysaetos*) chick banded at its nest in 2014 (MVZ:Bird:193216) with data from a radio transmitter device that tracked the individual’s last known location to Mexico in 2017; that observation was cataloged in Arctos as MVZObs:Bird:4792. Here, the coordinates for the original sampling locality are encumbered to protect the eagle nest. In another example (https://arctos.database.museum/guid/Arctos:Entity:134), a single endangered Mexican wolf (*Canis lupus baileyi*) was monitored through a federal conservation program with regular blood sampling at different places and times (e.g., MSB:Mamm:341613, MSB:Mamm:231704). Once moribund, the entire specimen was preserved and cataloged as MSB:Mamm:341614. Both examples illustrate how the Entity record functions to compile and unite multiple related occurrences or records of a single organism or collection object under one persistent identifier (Arctos base URL combined with the Darwin Core Triplet for the Entity record). The Entity identifier is passed to biodiversity data publishers via the Darwin Core organismID field, thus allowing aggregator portals and users to resolve these different records as the same individual. The flexibility of this model allows additional samples and observations to be continually linked to their Arctos Entity record as more data are collected.

### An educational resource

From its beginnings, Arctos has spearheaded collection-based inquiries for undergraduate education because of its web accessibility and richly linked data [3]. Students interested in biodiversity, evolutionary dynamics, spatiotemporal variation, cultural heritage, responses to anthropogenic change, and other topics have access to an array of data and tools that can initiate and answer interdisciplinary questions. This is exemplified by educational platforms where Arctos-based modules are posted for open-access class exercises (Table 4). Arctos also is used as a live classroom tool at universities (e.g., “Natural History Museums & Biodiversity Science” at the University of California Berkeley; “Museum Practicum in Advanced Collections Management” at the University of Colorado Boulder; “Mammalogy” at the University of New Mexico) and has been important in training undergraduate and graduate students, post-baccalaureates, and postdoctoral researchers in museum curation and data management [43]. Additionally, collection staff have used Arctos to creatively integrate museum objects with artwork and public engagement in an effort to educate students and the broader community about their collections. For example, the Alabama Museum of Natural History collaborated with the University of Alabama Fashion Archive to host a colorfest on social media that invited the public to interact with museum objects in art projects (#ColorOurCollections, http://library.nyam.org/colorourcollections; see Arctos project https://arctos.database.museum/project/10003310). In another art-based public exhibit, students, volunteers, and researchers spent a semester at the University of Wyoming Museum of Vertebrates creating original art pieces inspired by natural history objects, which were then displayed in a public show at the Berry Biodiversity Conservation Center (Art Inspired by Life, see Arctos project https://arctos.database.museum/project/10002855). At the University of Alaska Museum, staff in the Archaeology and Ethnology & History Departments work collaboratively with local, regional, national, and international organizations to highlight cultural items and Alaska Native heritage. As the Arctos network expands, so will its educational role in promoting awareness of the rich legacy and potential of museum collections.

**Table 4.**
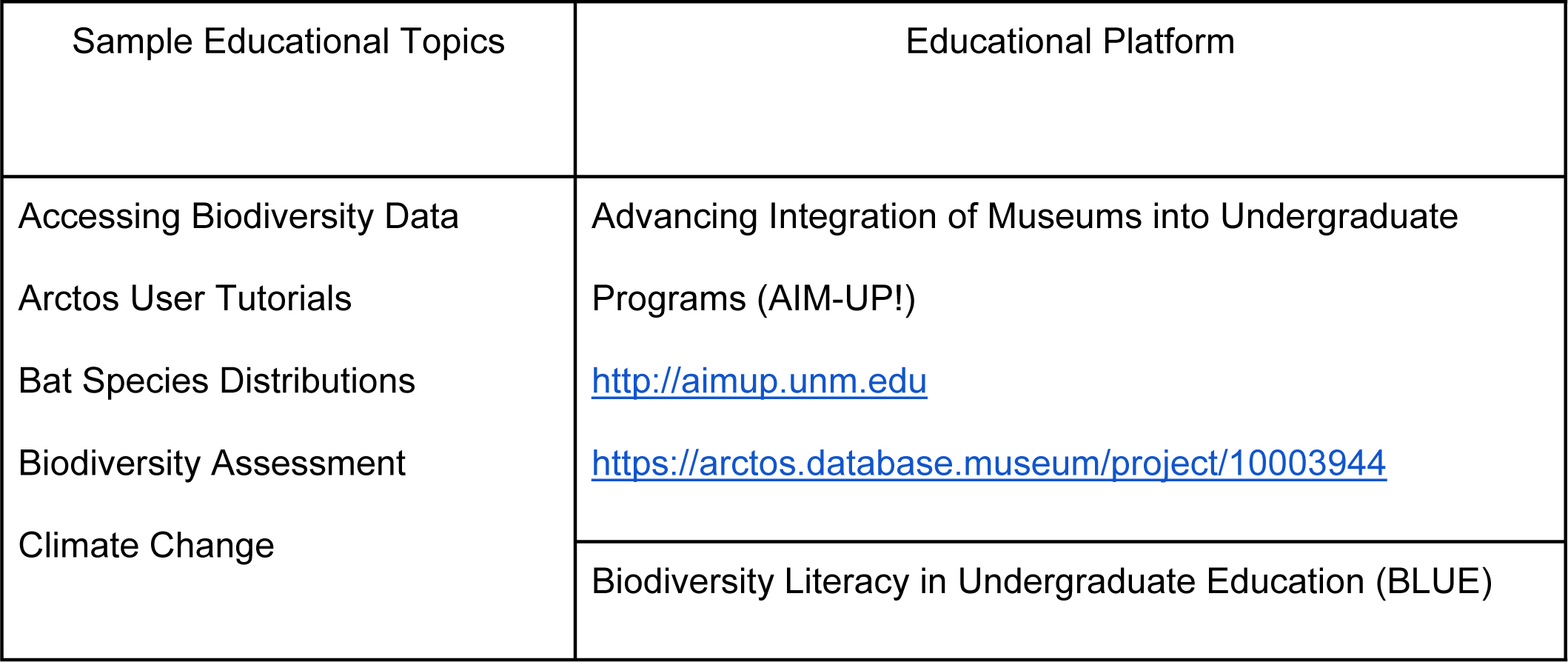
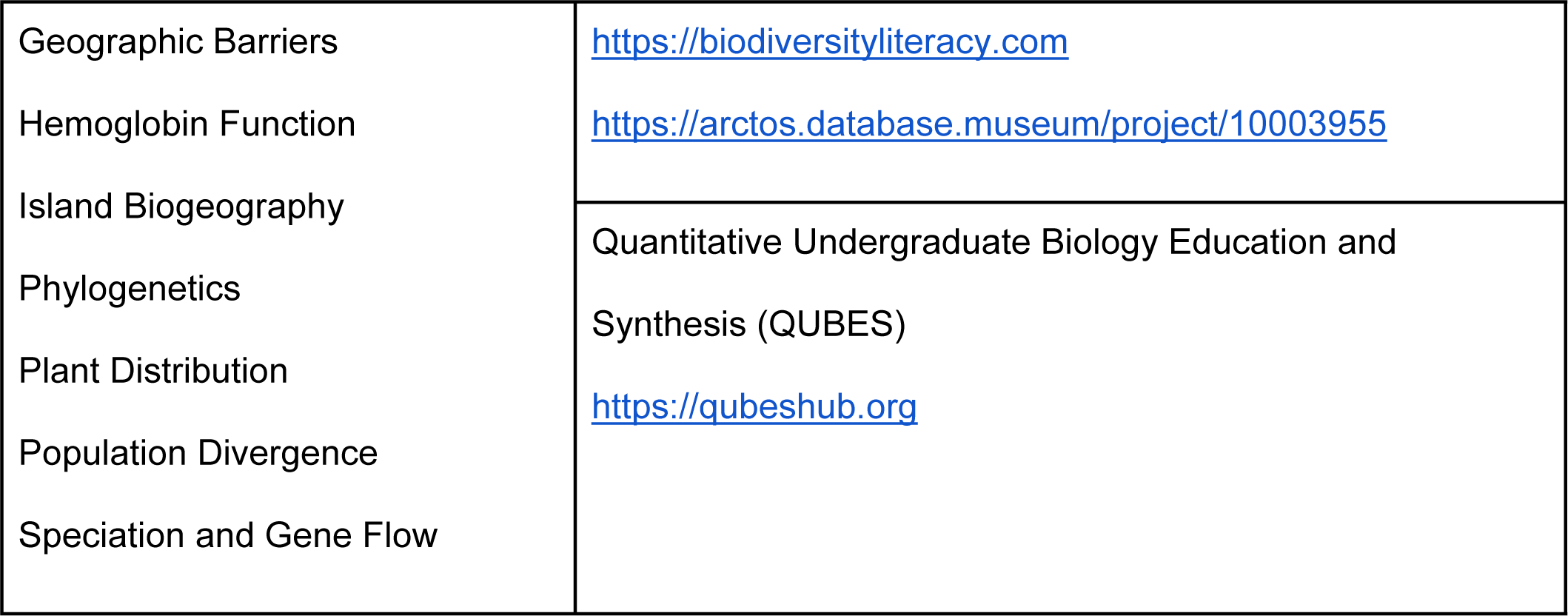
Examples of educational platforms with Arctos-based modules. Arctos modules aim to teach undergraduate students about biodiversity databases, biogeography, evolutionary biology, and climate change, among other topics.

### A sustainable future for Arctos

To fulfill the responsibilities of managing and ensuring access to our natural and cultural heritage data for current and future generations, museum administrators need to thoughtfully plan for the financial viability and health of their collections. Unfortunately, museum staff are too often overextended with diverse responsibilities and limited financial resources [44]. Another facet of sustainability is the maintenance and development of technical infrastructure. Arctos has a long track record as a model for “next-generation collections” and associated interdisciplinary research that addresses current and future societal challenges [17, 45–47]. Arctos’ longevity of nearly three decades may be due in part to its core development principles of standardization, flexibility, interdisciplinarity, and connectivity within a nimble development model for addressing novel needs and information types in response to changing technology, workflows, ethical considerations, and regulations [33, 48–50]. The sustainability and importance of maintaining these networked and interconnected technologies ultimately becomes premised on reliable funding. Despite the vital importance of these fundamental biodiversity digital resources, financial sustainability remains an ongoing community issue [51]. Facing this reality, the Arctos community sought to improve its financial model for the growing consortium of independent and diverse institutions. The most practical solution was fiscal sponsorship by a non-profit organization dedicated to supporting consortia like Arctos. This new business model, implemented in 2022, allows Arctos to follow diverse sources of funding and support including public and private grants, in-kind and volunteer assistance, fees for use, charitable donations, and annual subscription fees [44, 52]. Our overall goal is to use fiscal sponsorship to guarantee the success and sustainability of Arctos to ensure long term benefits to society and the community of biodiversity scientists, cultural heritage stewards, and educators of all kinds.

## Acknowledgements

We thank all members of the Arctos Working Group for their unflagging efforts to improve Arctos and keep it an active, functioning, and engaged community and platform. We also thank the generations of undergraduate and graduate students, post-baccalaureates, collection managers, curators, and technicians who perform daily collection tasks using Arctos at member institutions. The following individuals and collaborators have contributed invaluable expertise, perspectives, and support that have helped to enrich and expand Arctos as both a data platform and community: Stan Blum, John Deck, Jonathan Dunnum, Joyce Gross, Steffi Ickert-Bond, Gordon Jarrell, Craig Moritz, Kyndall B.P. Hildebrandt, Barbara Stein, Lam Voong, Cam Webb, John Wieczorek; Global Biotic Interactions (GloBI; Jorrit Poelen), Global Genome Biodiversity Network (GGBN; Katharine Barker), Integrated Digitized Biocollections (iDigBio; Gil Nelson, Deborah Paul, Erica Krimmel), and the Texas Advanced Computing Center (TACC; Chris Jordan). We thank the National Science Foundation for funding specific to the development and sustainability of Arctos (DBI-9630909, DBI-9876837, DEB-9981915, DBI-2034593, DBI-2034568, DBI-2034577), as well as the Robert & Patricia Switzer Foundation for awarding Arctos a Leadership Grant in 2023; additional grants from various sources have funded collection-specific initiatives that resulted in Arctos improvements. Finally, we thank Community Initiatives, especially Brandy Shah and Rose Cohen Westbrooke, for their guidance and expertise in our transition to fiscal sponsorship.

